# Community assessment of crustose calcifying red algae as coral recruitment substrates

**DOI:** 10.1101/2021.09.27.461929

**Authors:** Mari E. Deinhart, Matthew S. Mills, Tom Schils

**Affiliations:** Marine Laboratory, University of Guam, UOG Station, Mangilao, Guam; School of Science, Technology, and Engineering, University of the Sunshine Coast, Sippy Downs, Queensland, Australia

**Keywords:** coral larval settlement, coral reef restoration, CCA, Corallinophycidae, Corallinales, Peyssonneliales, Lithophylloideae, phylogenetics

## Abstract

Successful recruitment of invertebrate larvae to reef substrates is essential to the health of tropical coral reef ecosystems and their capacity to recover from disturbances. Crustose calcifying red algae (CCRA) have been identified as important recruitment substrates for scleractinian corals. As such, CCRA as a whole or subgroups (e.g., crustose coralline algae, CCA) are often used at the functional group level in experimental, ecological, and monitoring studies. Species of CCRA, however, differ in their ecological roles and their value as coral recruitment substrates. Here, we (1) investigate the species richness and community composition of CCRA on experimental coral recruitment tiles, and (2) assess if there is a recruitment preference of the coral *Acropora surculosa* for any of these CCRA species. 27 species of two orders of CCRA (Corallinales and Peyssonneliales) were identified from the recruit tiles. None of the DNA sequences of these species matched released sequences in GenBank or sequences of CCRA collected from natural reef systems in Guam. The similarity in CCRA communities between the recruitment tiles was high. Two species of CCRA were significantly preferred as recruitment substrates over the other CCRA species. Both of these species belonged to the subfamily of the Lithophylloideae. These two species are closely related to Pacific species that have been referred to as *Titanoderma* -but probably have to be assigned to another genus- and many of the latter have been attributed to be preferred coral recruitment substrates. Of all CCRA, Lithophylloideae sp. 1 had the highest benthic cover on the recruitment tiles and was the most preferred recruitment substrate. These findings highlight the high taxonomic diversity of CCRA communities and provide insight into species-specific ecological roles of CCRA that are often overlooked.

## Introduction

Coral reefs are threatened worldwide and are undergoing rapid change because of increased coral bleaching events, eutrophication, sedimentation, changes in land use and cover, and pest species outbreaks [1–6]. Apparent declines in live coral cover are generally the first effects that trigger concern for changes in benthic composition on coral reefs. Many other organisms are part of these intricate ecosystems but typically receive less scientific attention than scleractinian corals. Crustose calcifying red algae (CCRA; representatives of the red algal orders Corallinales, Sporolithales, Hapalidiales, and Peyssonneliales) are a dominant group of macroalgae on coral reefs that deposit calcium carbonate. Island groups in the tropical Pacific show significant differences in the abundance of CCRA on their reefs [7]. CCRA are essential components of healthy reef systems because of their ecological importance as (1) some of the main calcium carbonate depositors and carbon sequesters [8, 9], (2) the preferred settlement substrates for many invertebrate larvae (including scleractinian corals) [10], and (3) suppressors of nutrient indicator algae [11, 12]. Despite CCRA being among the most dominant benthic organisms in Guam’s tropical reef communities, CCRA systematics is still largely unexplored, consequently resulting in ecological gaps in knowledge.

CCRA contribute significantly to reef biodiversity [7, 8] and are a dominant component of Guam’s forereef community. When depositing calcium carbonate onto reefs in the form of high-magnesium calcite (Corallinophycidae) or aragonite (Peyssonneliales), CCRA cement and bind reef aggregates together, promote reef growth, provide protection against bioerosion, and enable the settlement of coral and other invertebrate larvae [11,13,14]. CCRA can also provide structural complexity and habitat diversity [15, 16].

The coral reefs of Micronesia are known for their high diversity of acroporid corals in the shallow forereef zones [17]. Acroporids are major reef builders due to their high benthic cover and fast growth rates [17, 18], thus creating habitat diversity for other reef organisms [19]. Guam’s acroporid corals have undergone extensive mortality in recent years, particularly in the forereef zone, due to bleaching events caused by episodes of elevated seawater temperatures in 2013, 2014, 2016, extreme low tides in 2015, and high predation from *Acanthaster planci* [1–6]. Understanding the factors that drive the recovery of reefs is important as the frequency, severity, and diversity of disturbances impacting coral reefs continues to increase [19]. The recovery of coral reefs depends on the regeneration of coral populations through the successful recruitment of coral larvae [20–22].

The continuing decline in acroporid [6] on Guam’s reefs has led to coral restoration efforts that primarily focus on rearing corals in nurseries for outplanting efforts (L. Raymundo, personal communication). Due to its overall resilience to disturbances in Guam, *Acropora surculosa* has been one of the main scleractinian coral species used for restoration efforts in Guam [6]. Corals used for restoration efforts are obtained through fragmentation of source colonies or via sexual reproduction. Sexually produced coral transplants can enhance genetic variability and can generate high numbers of new colonies to be used for large-scale restoration efforts [23, 24]. *A. surculosa* is a corymbose acroporid that has been well studied in Guam due to its occurrence in various reef habitats, its fast growth rate, and its overall value as a reef builder [25, 26]. *A. surculosa* is a hermaphroditic broadcast spawner, with spawning events occurring during the last quarter lunar cycles of July and August [27]. At the University of Guam Marine Laboratory, *A. surculosa* has been a popular study organism as part of master’s thesis projects [e.g., 28] and research publications [29–31].

CCRA have been well-documented to serve as the preferred settlement substrate for coral larvae [10,13,32–36]. Research has largely focused on the specific abiotic and biotic factors that facilitate successful scleractinian coral larval recruitment, settlement, growth, and fecundity [10,14,37]. The University of Guam Marine Laboratory has hosted *s*tudies describing various mechanistic pathways that facilitate *A. surculosa*, and *Leptastrea purpurea* larval settlement on CCRA [31,38, 39], yet it is unknown what CCRA species are favored for recruitment by *A. surculosa*. CCRA species can possess species-specific chemical fingerprints and microbiome communities [40], highlighting the need to correctly identify the CCRA that are favored for larval recruitment in the Micronesian region.

Experimental studies often identify CCRA byway of morpho-anatomical characteristics. Because of the requirement of specialized identification techniques [41–44] and extensive cryptic diversity in these red algae [45–50] identifications based on morphology and anatomy should be taken with reservations [50]. Species delineation of CCRA can be notoriously challenging due to their simple morphologies, their convergent evolution, their phenotypic plasticity that varies depending on habitat and life stage, and the frequent absence of reproductive features [45, 51]. Despite dispersal limitations, many CCRA species are reported to have broad geographical ranges based on morphological identification [52]. Overall, CCRA species richness of benthic communities is vastly underestimated [53, 54]. Due to these challenges, CCRA are generally treated as functional groups monitoring and ecological surveys, which does not recognize the different ecological roles that these CCRA serve on tropical reefs [54]. To address the challenges of CCRA species identification, DNA sequence analysis has proven to be an effective tool to investigate species diversity in CCRA [48, 49,55].

This study used DNA sequencing to investigate the species composition of CCRA communities on cured coral recruitment tiles in environmental conditions similar to natural reefs with pronounced *Acropora surculosa* stands (Pago Bay, Guam). Following the characterization of these CCRA communities, an analysis of recruitment preference by *A. surculosa* larvae for CCRA species was conducted. We hypothesized that (1) a diversity assessment using DNA barcoding would reveal more CCRA taxa than a morphological diversity assessment and (2) *A. surculosa* larvae prefer to recruit on a select number of CCRA taxa. Based on previous recruitment studies [10,34,35] we hypothesize that *A. surculosa* larvae will favor members of the subfamily Lithophylloideae (Corallinales, Rhodophyta) as recruitment substrates. This study aims to provide new insights on CCRA diversity and community composition in the Western Pacific and the significant ecological roles fulfilled by CCRA species.

## Materials & methods

### CCRA species on coral recruitment tiles

Coral larval recruitment and settlement in a controlled environment can protect corals from harsh conditions during their most vulnerable stages [56]. The Raymundo Coral Lab follows SECORE protocols for rearing and curing coral recruitment tiles in the flow-through seawater system of the University of Guam Marine Laboratory. Seawater is drawn from a coral reef in Pago Bay on the eastern shore of Guam. Recruitment tiles were cured in holding tanks of a seawater flow-through system to have them covered with CCRA before the *A. surculosa* spawning event in July 2018. Twelve coral recruitment tiles were cured in the same tank, subsequently there was little to no variation in temperature, salinity, sediment load, and water depth between the tiles.

Twelve star-shaped coral recruitment tiles with successful coral settlement following the 2018 *A. surculosa* spawning event were used for this study. The eleven exposed sides of each tile were labeled and photographed before CCRA tissue samples were scraped off for DNA extraction. All CCRA on which coral recruits were detected were sampled for DNA extraction. In addition, 32 samples of different CCRA morphologies that were not associated with coral recruits were also collected. Samples without associated coral recruits were chosen based on their unique morphology and replicate samples of the different morphologies were taken to address cryptic diversity.

### DNA barcoding

A total of 92 CCRA specimens were identified using DNA sequencing. For each specimen, a patch of tissue free from epiphytes was swabbed clean with a 10% bleach solution. A Dremel rotary tool, a pair of tweezers, or single-edged razor blades were used to scrape off tissue from each specimen for DNA extraction. The Dremel, tweezers, and razor blades were sterilized by soaking them in 10% bleach and heating them over a flame after each tissue removal to avoid contamination. Tissue samples were placed in sterile 1.5 mL Eppendorf tubes for DNA extraction. DNA of each algal specimen was extracted using QIAGEN DNeasy Blood & Tissue Kit (Qiagen Inc., Valencia, CA) or the GenCatch Blood & Tissue Genomic Mini Prep Kit (Epoch Life Science Inc., Missouri City, TX) following the manufacturer’s bench protocol.

Three genetic markers were amplified via polymerase chain reaction (PCR) for species delimitation and identification. The mitochondrial cytochrome c oxidase subunit 1 DNA barcode region, COI-5P (roughly 664 bp), allows for species delimitation of CCRA and it is the official barcode marker for DNA barcoding of red algae [45]. The primer combination used to amplify COI-5P was TS_COI_F01_10 (5’-TCGARTCYCGTCTCTCTCG-3’) [57] and the reverse primer, GWSRx [58]. Protocols developed by Mills & Schils [57] were followed for COI-5P amplification.

Chloroplast photosystem II thylakoid membrane protein D1, *psb*A (roughly 950 bp), was a second marker used for DNA barcoding and species delimitation in the Corallinales. The *psb*A marker is more conserved than COI-5P and is frequently used for CCRA identification because of its high success rate of amplification. The primers used to amplify this gene were psbAF and psbAR2 [59]. Amplification of *psb*A followed the PCR protocol outlined by Mills & Schils [57]. The chloroplast ribulose-1, 5-biphosphate carboxylase large subunit, *rbc*L (roughly 1,350 bp), was amplified for a subset of Corallinales specimens from the coral recruitment tiles. Amplification of *rbc*L used the primers F57 and rbcLrevNEW following the amplification profile [60].

### Species delimitation and phylogenetic analysis

PCR products were sent to Macrogen Inc. (Seoul, Republic of Korea) for DNA sequencing. Once chromatograms were obtained, consensus sequences were assembled using Geneious Pro 11.0.5 computer software (https://www.geneious.com; [61]). Consensus sequences where then compared with available CCRA sequences via BLAST search (Basic local alignment search tool) [62] or available CCRA sequences from online repositories such as GenBank and the Barcode of Life Database (BOLD) [63]. Sequences of previously collected CCRA samples from Guam were also compared to the sequences of the coral recruitment tiles for species delimitation. Gene alignments were created using the MUSCLE plugin [64] in Geneious Pro 11.0.5. An alignment of COI-5P was made for the Peyssonneliales and a concatenated alignment of COI-5P, *psb*A, and *rbc*L for the Corallinales. To assess species richness and delimitate putative species in this study sequence divergence analyses were calculated using the Automatic Barcode Gap Discovery (ABGD) [65] for COI-5P (<3%), *psb*A (<2.5%), and *rbc*L (<0.9%) [56].

To resolve the taxonomic identity of CCRA, all sequences from the coral recruitment tiles were selected for phylogenetic analysis. Maximum likelihood (ML) was used to infer phylogenies through IQ-TREE 2 [66]. IQ-TREE 2 uses a combination of hill-climbing approaches and stochastic NNI operations to obtain higher likelihoods while estimating maximum likelihood phylogenies [66]. The concatenated alignment was partitioned according to gene and the optimum evolutionary model for each gene was found using ModelFinder [67]. The Ultrafast Bootstrap Approximation was used to achieve unbiased node support values with 1000 replicates [68].

Corallinales species could not be identified to family or genus level after BLAST searches [62]. To identify Corallinales specimens, their sequences were aligned and analyzed against the seven-gene concatenated alignment of Peña et al. [69] (S1 Fig). This alignment of Corallinophycidae (S1 Table; S2 Table; S1 Fig) consisted of seven genes: 23S rRNA, COI, EF2, LSU rRNA, *psb*A, *rbc*L, and SSU rRNA. The total length of the seven-gene concatenated alignment was 11,608 bp. The final length of each alignment resulted in: 370 bp for 23S rRNA, 687 bp for COI, 1,622 bp for EF2, 4,716 bp for LSU, 784 bp for *psb*A, 1,386 bp for *rbc*L, and 2,086 bp for SSU. ModelFinder [67] in IQ-TREE 2 [66] found that the best-fit partition model for each partition were: TVMe+I+G4 (23S rRNA), TN+F+I+G4 (EF2), TIM3+F+I+G4 (LSU), and GTR+F+I+G4 (COI, *psb*A, *rbc*L, and SSU).

BLAST searches could not identify the 10 of the putative Peyssonneliales taxa to genus level. A separate phylogenetic analysis was run for Peyssonneliales taxa of the recruitment tiles to identify taxa to the lowest taxonomic level possible. The 32 Peyssonneliales sequences from this study were aligned with 11 Peyssonneliales COI-5Psequences (S3 Table) with *Bonnemaisonia asparagoides* (Woodward) C. Agardh as the outgroup. Priority of comparison sequences was given to type species of each genus. If type species sequences were unavailable, sequences of congeners were used. The available sequences belonged to the genera *Incendia, Metapeyssonnelia, Peyssonnelia*, *Polystrata*, *Ramicrusta*, *Riquetophycus*, and *Sonderophycus* to help resolve taxonomic identification of the putative species, with *Bonnemaisonia asparagoides* (Bonnemaisoniales) as the outgroup. The length of the COI-5P Peyssonneliales alignment was 621 bp. A maximum likelihood phylogeny was generated through IQ-TREE [66]. The best-fit evolutionary model was GTR+F+I+G4 (ModelFinder) [67].

### CCRA and substrate cover on coral recruitment tiles

Nine substrate categories were visually discerned on the coral recruitment tiles. The identity of these substrate categories was investigated and validated using DNA sequence analysis. Grouping CCRA taxa into substrate categories based on their visual recognition was required for the benthic cover and settlement preference analyses (Table 1). Photographs of each tile and its 11 sides were taken on 13 December 2018. Substrate cover was measured using Adobe Photoshop version CC 2020 software by converting the total pixel counts to surface area measurements for each tile side. Color overlays were used to obtain the pixel count for each substrate category present on the tiles. The benthic cover of each category was calculated by dividing the total number of pixels of the color overlay for each category by the total pixel count of the tile side (Fig 1). Pixels of coral recruits were ascribed to the substrate category on which they recruited.

**Fig 1.**
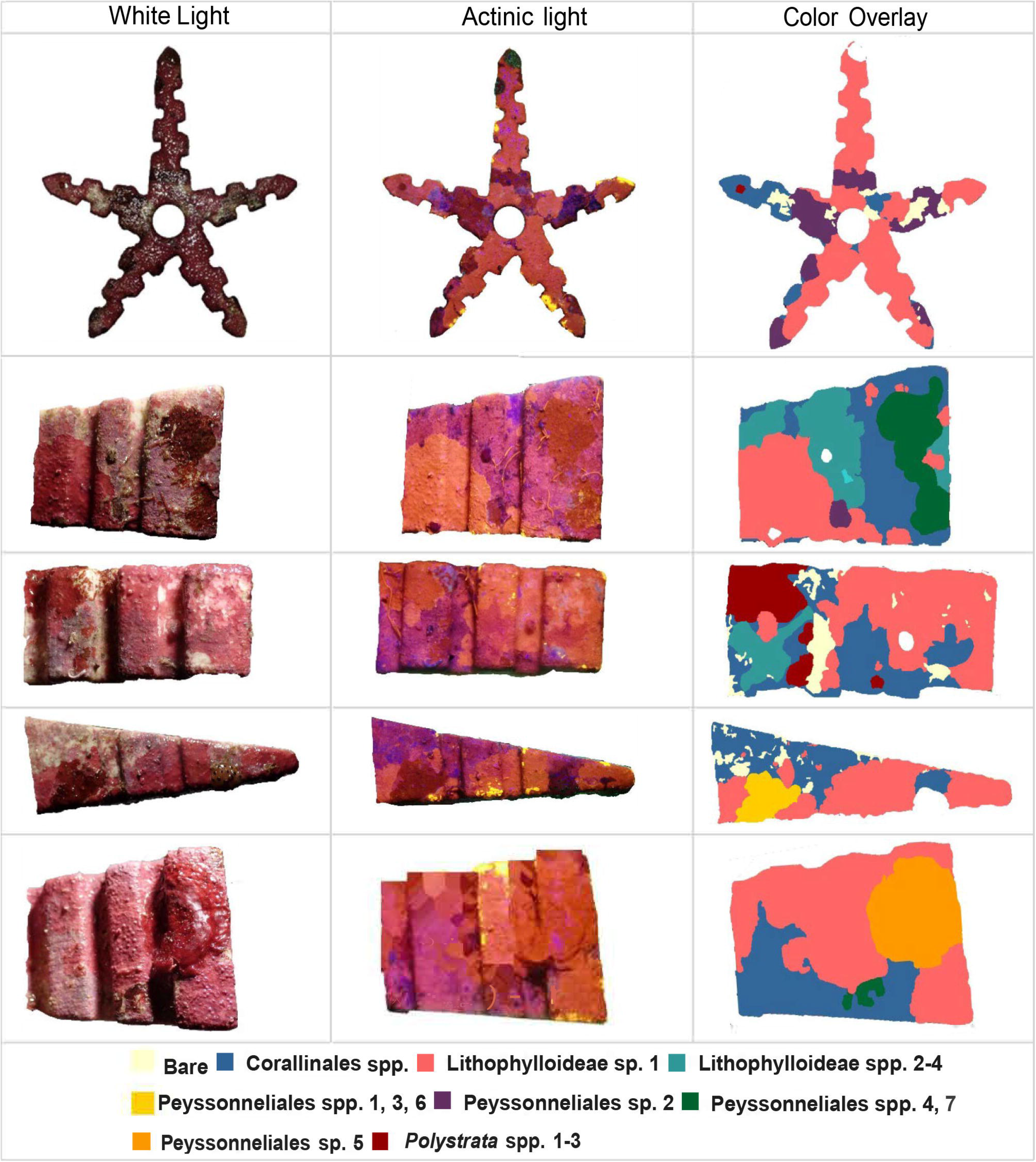
Example of coral recruitment tile photographs under white and actinic light plus the color overlay used for the calculation of percent cover for each substrate category. Benthic cover was assessed for all 11 exposed sides of the 12 tiles. Substrate category colors are similar to those in Figs 4, 5.

**Table 1.**
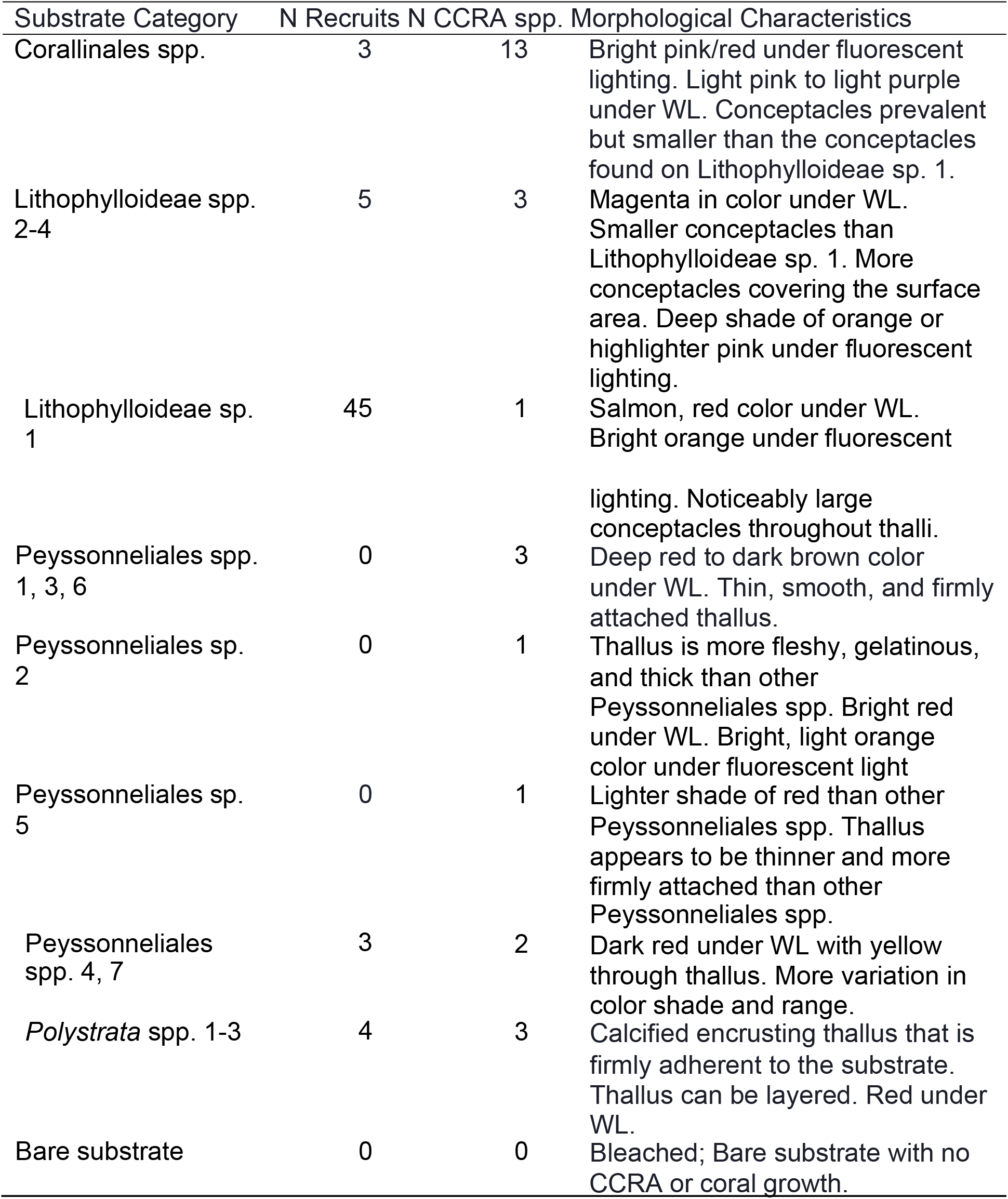
The nine substrate categories used for image analysis and the number of coral recruits per substrate category. Colors correspond with the category they were placed in. The “species count” column indicates how many species were placed in each substrate category. The “morphological characteristics” column provides a description of the features that could be visually distinguished for each substrate category during the image analysis.

### Statistical analysis

Data analysis was carried out in the R environment for statistical computing and visualization (v 3.5.1; R Development Core Team 2020). The percent cover of substrate categories on recruitment tiles was presented as mean ± standard deviation. A Kruskal-Wallis H test analyzed to see if the cover of CCRA substrate categories were similar to each other or if there were differences in cover between the substrate categories, followed by Tukey’s Honest Significant Difference test as a post-hoc test. The Tukey test compared the mean cover of each substrate category to the means of all substrate categories. If a substrate showed to have a significantly different cover than the other substrates, another Kruskal-Wallis test was conducted to identify if there was any significant difference in cover of the dominant CCRA substrate between the 12 tiles.

The G-test for goodness-of-fit (likelihood ratio or log-likelihood ratio) was used to evaluate whether *Acropora surculosa* larvae (1) were evenly distributed over the recruitment tiles and (2) preferred to recruit on specific substrate categories over others. This G-test was chosen because of the small sample size and the existence of one nominal variable (coral recruitment) with more than two values (substrate categories). The G-test evaluates if the observed number of coral recruits differs significantly from the expected number of recruits on individual tiles or on each of the substrate categories based on their percent cover. The null hypothesis for tile recruitment preference was that their occurrence was evenly spread over all the tiles. The null hypothesis for substrate recruitment preference was that the number of coral recruits per substrate category was a direct function of the percent cover of each substrate category, *i.e.* random settlement.

## Results

### Species delimitation

The 91 CCRA samples extracted from the coral recruitment tiles resulted in 84 successful DNA sequences, all belonging to the class Florideophyceae. Of the 84 successfully sequenced samples, 52 samples belonged to the order Corallinales, while 32 samples belonged to the order Peyssonneliales. The 84 samples represented 27 distinct species (Table 1). Of these 27 species, 17 species belong to the order Corallinales and 10 belong to the order Peyssonneliales. The amplification of all three genetic markers for each putative species was not always successful, however 16 Corallinales species had a successful amplification for at least two markers. No sequences from GenBank or BOLD matched (>97% sequence similarity) the 27 CCRA species from the coral recruitment tiles.

Five samples could not be sequenced but were visually identical to specimens that were successfully sequenced. One of these samples was identified as Lithophylloideae sp. 1, due to its similarity in habit to the 33 successfully sequenced samples of this species and one was identified as Lithophylloideae sp. 2–4. Two other samples were confidently identified as Peyssonneliales spp. One remaining CCRA corresponded to other Corallinales taxa that were not identified to a subfamily level.

### Corallinales

Out of the 17 Corallinales species, 16 were identified as members of the family Lithophyllaceae with representatives for each of the four subfamilies: Lithophylloideae, Hydrolithoideae, Chamberlainoideae, and Metagoniolithoideae (S1 Fig). One Corallinales species was identified as a representative of Neogoniolithoideae (Corallinaceae, Rhodophyta).

Seven of the putative Lithophyllaceae species are members of the subfamily Lithophylloideae (Fig 4). Lithophylloideae sp. 3 and Lithophylloideae sp. 4 are sister taxa to *Lithophyllum pustulatum* (J.V. Lamouroux) Nägeli and two other *Titanoderma* spp. in SI Fig [69]. The validity of the genus *Titanoderma* is being debated [50, 70] and therefore the species from the recruitment tiles are referred to as Lithophylloideae spp. Lithophylloideae sp. 1 and Lithophylloideae sp. 2 are sister taxa to species named *Titanoderma* sp. and *Lithothrix* sp. in Peña et al. [69] (S1 Fig). However, the type species of *Titanoderma* (*T. pustulatum*) was resolved in a disparate clade (S1 Fig). Therefore, Lithophylloideae spp. 1 and 2 are not *Titanoderma* species. Lithophylloideae sp. 1 was represented by the most sequences in this study. Further phylogenetic analyses are required to resolve the taxonomy of these species and they are provisionally named as distinct species within the subfamily.

Five Corallinales species were confidently assigned to genus level. One species belonged to the genus *Hydrolithon* (Hydrolithoideae, Rhodophyta) with high support values in the phylogeny (Fig 2). Four CCRA species belonged to the subfamily Metagoniolithoideae (Fig 2): *Harveylithon* sp. 1, *Harveylithon* sp. 2, *Harveylithon* sp. 3, and *Porolithon* sp. 1. All four species grouped with their congeners with high support values. *Harveylithon* sp. 2 and *Harveylithon* sp. 3 are sister taxa to each other, while *Harveylithon* sp. 1 is a sister taxon of a *Harveylithon* sp. from Vanuatu [69] (S1 Fig). *Neogoniolithon* sp. 1 was the only species in this study that represented the family Corallinaceae (Fig 2). Support values for the genus *Neogoniolithon* were high (>100), which included *Neogoniolithon* sp. 1 (S1 Fig).

**Fig 2.**
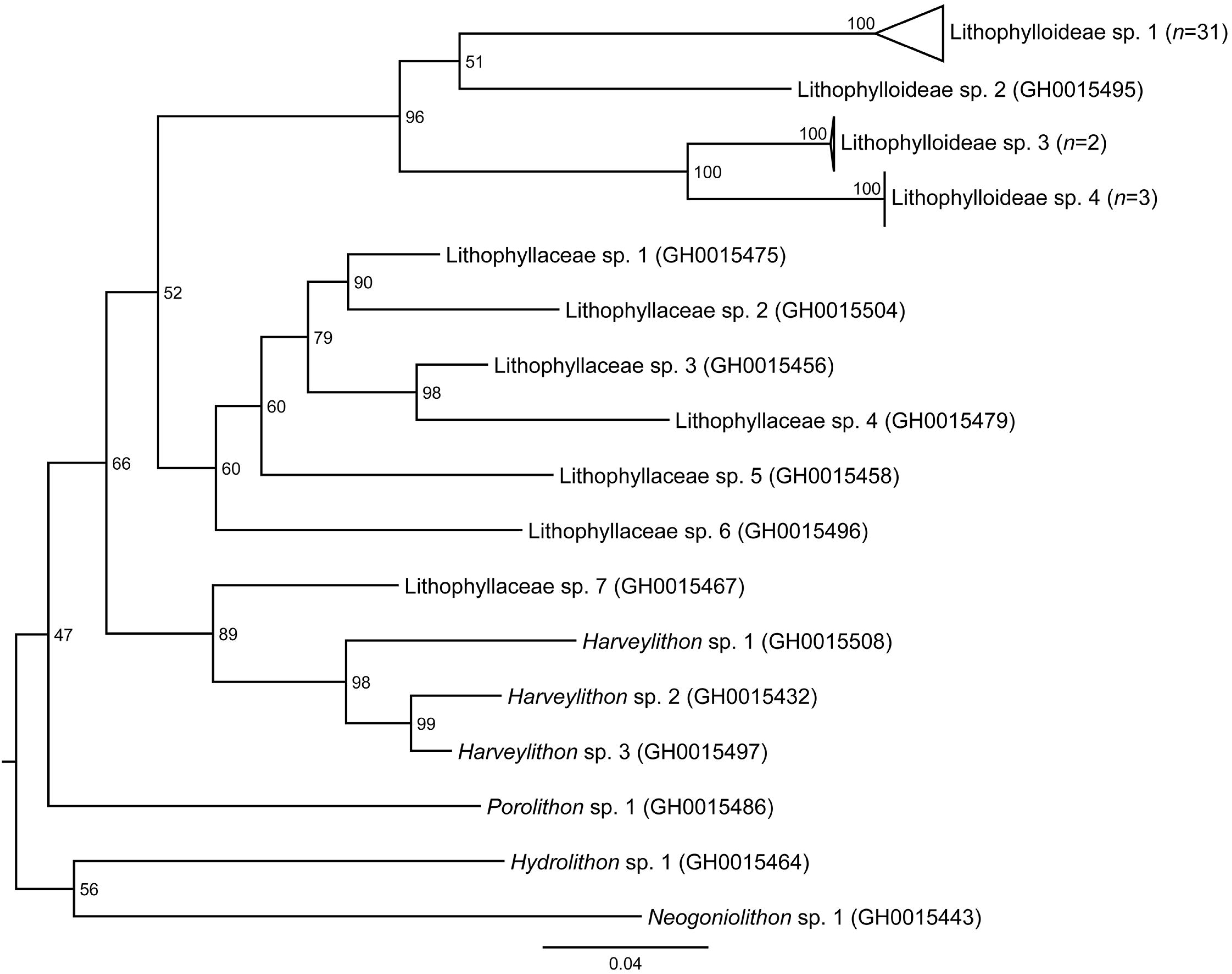
Maximum likelihood phylogeny of the 17 Corallinales species from the recruitment tiles inferred from COI-5P, *psb*A, and *rbc*L. GH numbers are the specimens’ voucher numbers in the Guam Herbarium (GUAM). The tree was inferred by maximum likelihood with IQ-TREE [66]. Support values listed next to each node are ultrafast bootstrap approximations based on 1000 replicates. Scale bar represents nucleotide substitutions per site.

Seven Corallinales species could not be assigned to a subfamily (Fig 2). Lithophyllaceae spp. 1-6 could not be placed readily into a recognized genus or subfamily (S1 Fig). These six species form a clade within the Lithophyllaceae and sister to the Metagoniolithoideae (Fig 2). Therefore, these taxa were referred to as distinct species within the family Lithophyllaceae, i.e. Lithophyllaceae spp. 1–6. Each of these Lithophyllaceae spp. was represented by just a single sample, with at least 2 successfully sequenced markers. Lithophyllaceae sp. 7 also did not group with any other clade. The nearest clade to Lithophyllaceae sp. 7 was the subfamily Hydrolithoideae and the recently described genus *Parvicellularium* [71] (S1 Fig). Lithophyllaceae sp. 7 was a singleton specimen with only a successful *psb*A sequence.

### Peyssonneliales

COI-5P sequences of Peyssonneliales taxa discerned 10 species on the coral recruitment tiles. Three of these species were representatives of the genus *Polystrata* and seven species belonged to two distinct clades that could not be assigned to a recognized genus within Peyssonneliales (Fig 3). Peyssonneliales spp. 1–7 did not match nor were closely related to any sequenced species through a BLAST search. These seven Peyssonneliales members are not part of the clade with the type species of the genus *Peyssonnelia*, *Peyssonnelia squamaria* Gmelin. Peyssonneliales sp. 1 and Peyssonneliales sp. 2 form a strongly supported clade. Peyssonneliales spp. 3–7 were resolved in another clade, with strong support for the subclade consisting of Peyssonneliales spp. 3–6 (Fig 3). Three *Polystrata* species were identified on the coral recruitment tiles and clustered together as a monophyletic genus (Fig 3).

**Fig 3.**
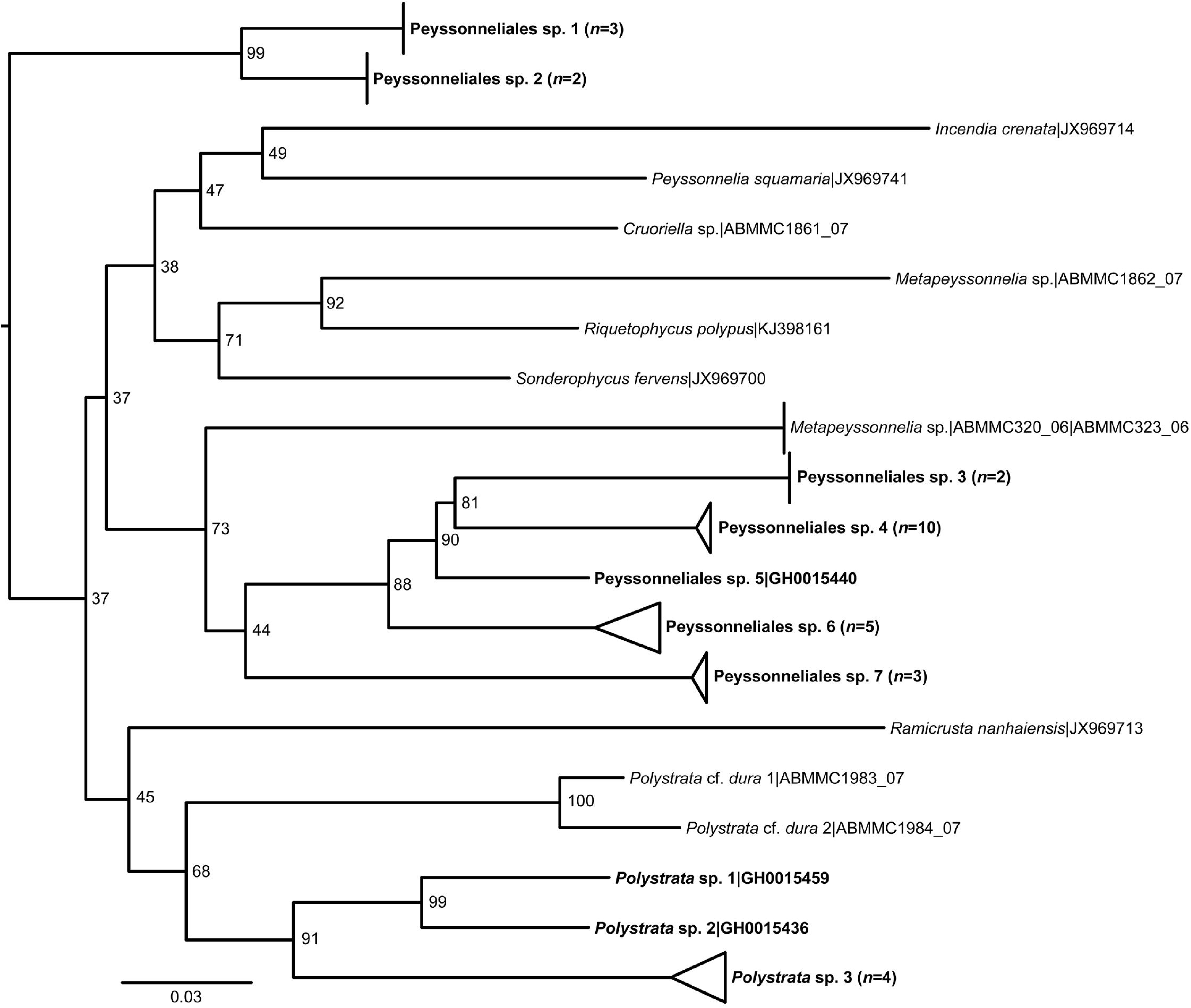
Maximum likelihood phylogeny of the 10 Peyssonneliales species (boldface font) from the recruitment tiles and 10 reference sequences from GenBank inferred from COI-5P sequences. The tree was inferred by maximum likelihood with IQ-TREE [66]. Support values listed next to each node are ultrafast bootstrap approximations based on 1000 replicates. Scale bar represents nucleotide substitutions per site.

### Substrate Composition and Acropora surculosa Settlement Preference

The 27 CCRA species were grouped into ten substrate categories based on their phylogenetic placement and habit similarities (Fig 2, 3; Table 1). The cover of all twelve tiles was comparable and dominated by healthy CCRA (Fig 5). Corallinales members constituted the highest percent cover on the tiles (>50%), followed by bare substrate and stressed CCRA (∼25%), and Peyssonneliales constituted the lowest cover (<20%; Fig 4). Although the Corallinales communities were more species rich than those of the Peyssonneliales, Corallinales species were more difficult to discern visually compared to Peyssonneliales species.

**Fig 4.**
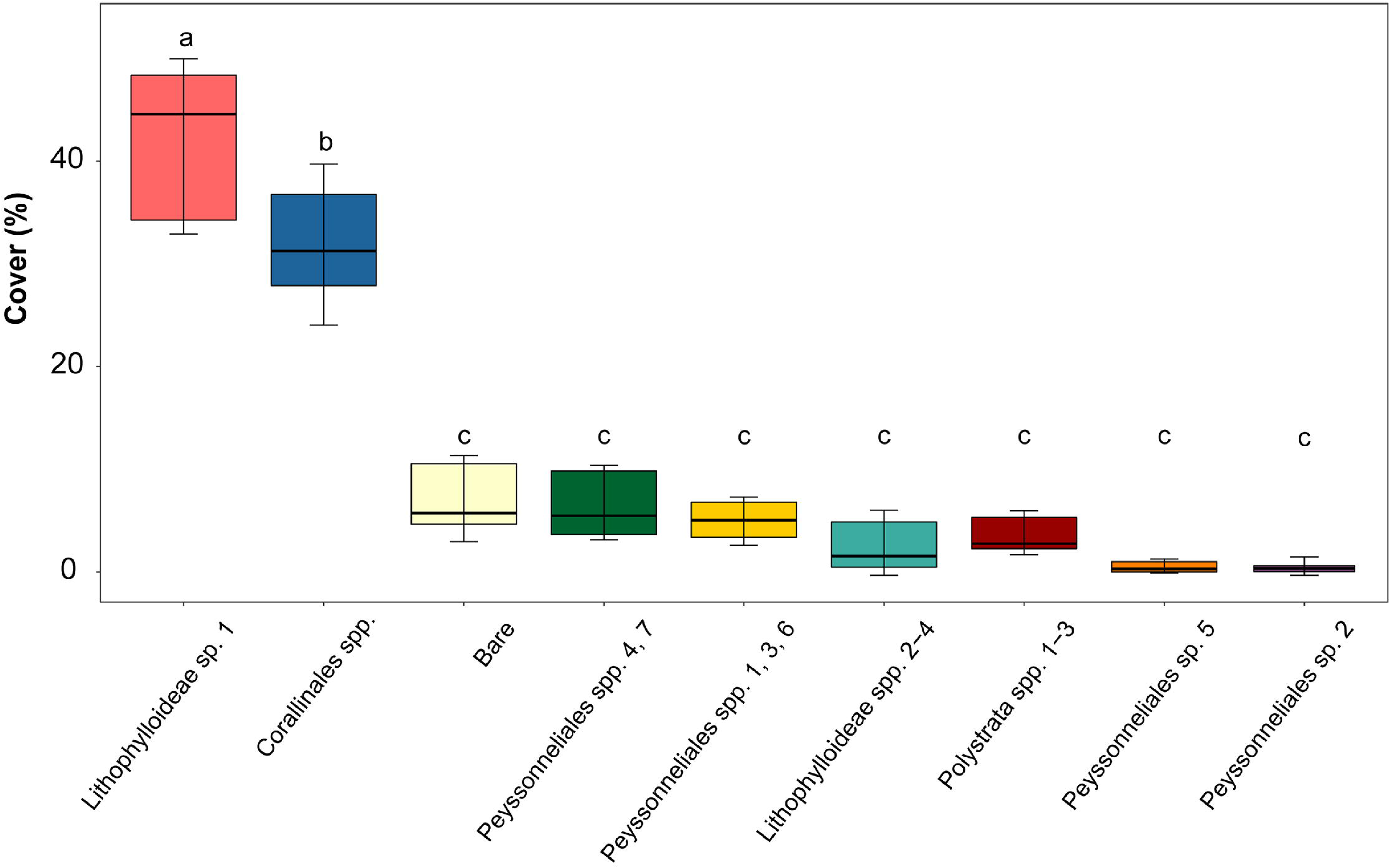
Box plot of the percent cover for each substrate category on the 12 coral recruitment tiles. The abscissa lists substrate categories, the ordinate shows percent cover. Median cover of a substrate category is indicated by a horizontal line in the interior of the box. The box represents the inter-quartile range (IQR) between the upper and lower quartiles. Whiskers indicate the minimum and maximum values beyond the IQR. Letters indicate non-significant differences between island groups (*P*>0.05). Box colors match substrate category colors in Figs 1, 5.

**Fig 5.**
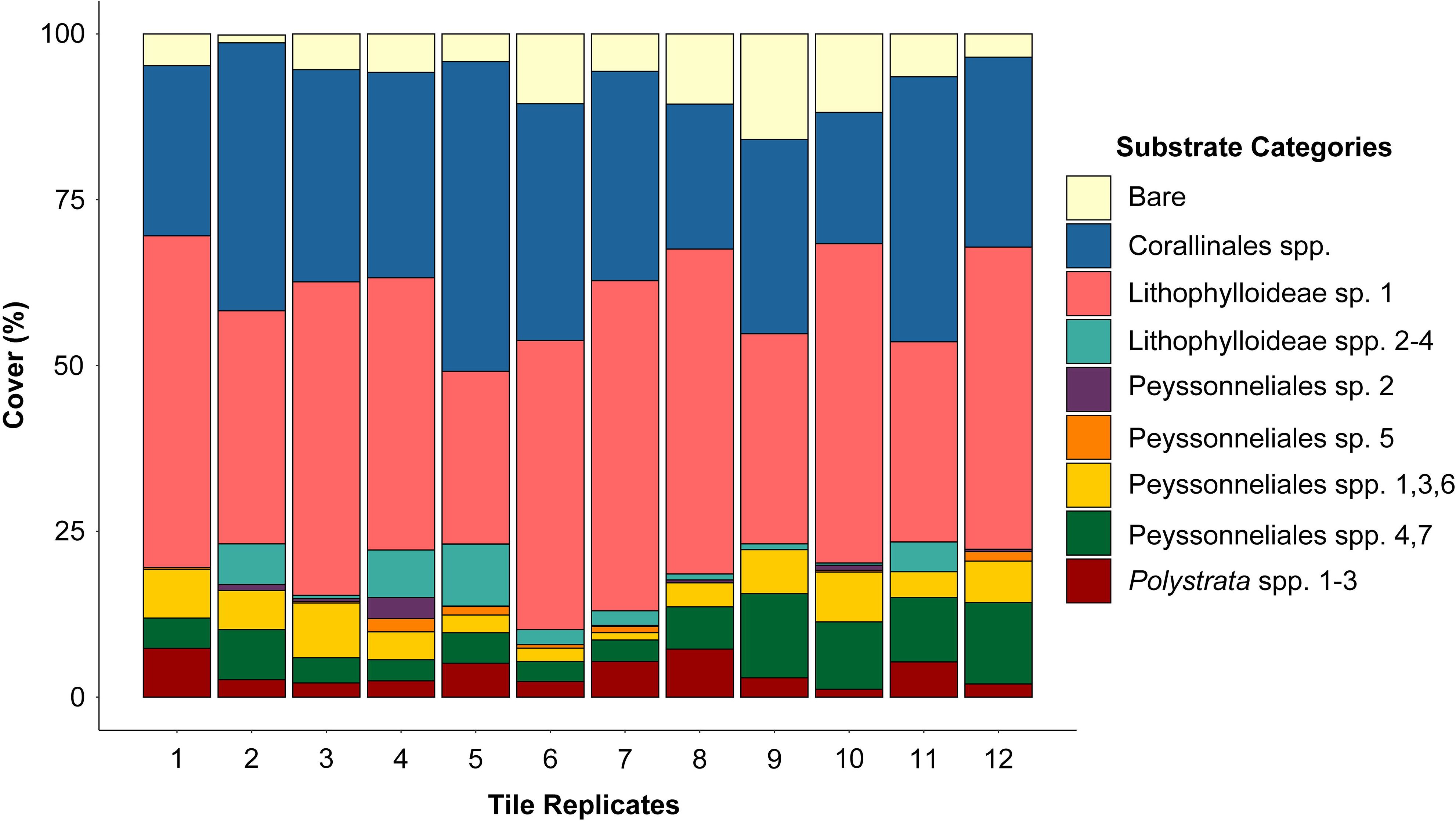
Substrate community composition of the 12 coral recruitment tiles. The abscissa lists the 12 recruitment tiles, the ordinate shows percent cover. Colors match substrate category colors in Figs 1, 4.

During the image analysis for substrate measurement, it was noted that four coral recruits had been overlooked during the CCRA sampling process, due to their small size. These coral recruits were identified in the digital images and included in the analyses with the substrate category that they recruited to (*i.e.*, Corallinales spp. and Lithophylloideae sp. 1). All coral recruits were associated with CCRA. Just one coral recruit was associated with a Corallinales spp. with an unhealthy appearance (loss of pigmentation, but not bleached).

The Kruskal-Wallis H test resulted in showing there was a significant difference in cover between the substrate categories and there was a substrate category that was significantly dominant on the tile community (p = 3.961e-13). Tukey’s post hoc test concluded that the nine substrate categories fell in one of three groups based on their percent cover across the tile community and identified Lithophylloideae sp. 1 to be the dominant substrate category (Fig 4). Substrate composition of recruitment tiles was similar between tiles (Fig 5). The overall cover of Lithophylloideae sp. 1 across the 12 tiles was similar, meeting the Kruskal-Wallis H test null hypothesis (*p* = 0.4433). The number of coral recruits on each of the twelve tiles did not differ significantly from random recruitment. The G-Test for goodness of fit also showed that *A. surculosa* larval settlement was not random and that certain substrate categories were favored for recruitment (*p* < 0.0001). Each substrate category was then individually tested for recruitment preference using the G-test. A total of 33 samples of Lithophylloideae sp. 1 were successfully sequenced from the coral recruitment tiles, which was consistently the dominant species (41.5%) on each tile (Fig 4). The habit of Lithophylloideae sp. 1 could confidently be recognized on the coral recruitment tiles and in the photo analysis to reliably calculate its percent cover. Of the 60 coral larvae present on the tiles, 45 of them recruited on Lithophylloideae sp. 1 (Table 1). Several of the sequenced Lithophylloideae sp. 1 samples had more than one coral recruit associated to them. Because Lithophylloideae sp. 1 was the dominant substrate on the tiles, the G-test clarified that the higher number of coral recruits on this species was more than just the result of its high percent cover. *A. surculosa* larvae significantly preferred this species as a recruitment substrate (*p* = 1.1308e-07).

Lithophylloideae spp. 2–4 was the substrate category with the second most coral recruits (5; Table 1). The average surface area of Lithophylloideae spp. 2–4 was 2.9% per tile (Fig 4). Despite its low cover, *A. surculosa* larvae also significantly preferred to recruit on this substrate category (*p* = 0.0373). Out of the seven remaining substrate categories, three substrates had coral recruits on them: Corallinales spp., Peyssonneliales spp. 4, 7 and *Polystrata* spp. 1–3. Corallinales spp. was the category that comprised the highest number of species (13 spp.; Table 1). Many of the species were represented by a single sample. Corallinales spp. comprised a large percent cover on the recruitment tiles (31.9%; Fig 4). Coral settlement to Corallinales spp. was significantly less than expected based on its percent cover on the tiles (*p* = 2.273e-07). The remaining two categories with coral recruits were not statistically significantly preferred as recruitment substrates: Peyssonneliales spp. 4 and 7 had four recruits (6.8% cover; *p* = 0.977) and *Polystrata* spp. 1–3 had three recruits (3.8% cover; *p* = 0.6501).

## Discussion

Successful dispersal and recruitment of invertebrate larvae are crucial to the health and recovery of tropical reef ecosystems following environmental disturbance [72]. Prior to this study, it was unclear which CCRA species served as the preferred recruitment substrates of *A. surculosa* in the western Pacific. This study reports on a diverse CCRA community composition on newly colonized bare substrates in a semi-natural environment, and the CCRA species that are significantly favored by *A. surculosa* recruits.

### CCRA species diversity and community composition

CCRA are a speciose and phylogenetically diverse group that is often treated as a single functional group in ecological and experimental studies. Studies on the taxonomic and ecological composition of CCRA communities are scarce [73] and are typically based on morphological observations [34,74,75]. By tissue sampling in conjunction with tile imaging, we were able to describe the CCRA community that settled on the coral recruitment tiles by linking molecularly assigned species with habit observations. This allowed us to calculate the cover of closely related CCRA species on coral recruitment tiles. The CCRA species on the recruitment tiles in the flow-through seawater system are probably a subset of the CCRA flora on the shallow forereef zone of Pago Bay where the seawater is drawn from.

Molecular diversity studies of CCRA have consistently surpassed species richness estimates based on morphological analyses [48,76–78]. The most recent account of CCRA diversity for Guam was summarized in a checklist of Guam’s seaweed flora [79], which lists 24 CCRA species for Guam. Of these 24 species, 17 belong to the Corallinales, two belong to the Peyssonneliales, four belonging to the Hapalidiales, and one belongs to the Sporolithales. Recently, two new species records and four new species of the genus *Ramicrusta* (Peyssonneliales) were reported for Guam through molecular-assisted alpha taxonomy [57], increasing Guam’s CCRA species count to 30. All of the 27 species on the recruitment tiles are representatives of the Corallinales and Peyssonneliales and hereby doubling the species richness of these orders for Guam [79]. The species richness exceeded what could be observed using only anatomical observation. We do note that only one sample for each of the 13 Corallinales species that were not members of the Lithophylloideae subfamily were sequenced, leading to challenges in morphologically distinguishing each species from the other during the measurement assessment. Had DNA sequencing not been utilized for this study, CCRA diversity on the tiles would have been limited. These results provide further support for a significantly higher CCRA diversity in Guam than what is currently reported.

Advances in molecular techniques have increased the scale of DNA barcoding efforts in floristic studies, which has allowed for more accurate and rapid diversity assessments than studies based on morpho-anatomical identifications. For example, 72 new records of algal species, including members of Corallinales and Peyssonneliales, were reported for four shallow reefs sites in northern Madagascar [78]. Similarly, DNA barcoding identified 122 CCRA species for various sites in southern New Zealand with the recognition of CCRA diversity hot spots [48]. Considering the small surface area from which our 92 CCRA samples were collected (106 cm^2^ per tile), which further supports the notion that CCRA species richness at small spatial scales can be hyper-diverse, particularly in areas with high microhabitat diversity.

Community composition between the tiles was similar with high CCRA richness per tile. Most substrate categories were found on all tiles and the proportional abundance of substrate categories was similar between tiles. Lithophylloideae sp. 1 was the dominant species on all 12 tiles (41% ± 8.53 SD), and other substrate categories were significantly less abundant. Algal turfs constitute similar communities as CCRA composed of a patchwork of small algae. These turf communities have also been characterized by high species diversity [80, 81]. Turf algal communities in Lhaviyani Atoll, Maldives, were found to be species rich and highly variable at small spatial scales (samples were separated by 10 cm) [82]. The CCRA communities on recruitment tiles contained a similarly high species richness but were homogeneous in composition between tiles. This limited diversity between tiles probably results from the replicate study design in which environmental and microhabitat diversity was minimized.

### Identification of CCRA taxa

Molecular studies have become the norm for species delineation in phycology, resulting in the detection of hundreds of new species at a rapid pace [83]. While this has broadened our understanding of algal diversity, it has created challenges to describe and name algal species, including CCRA, since formal descriptions are time-consuming and trail behind the molecular identification of species [44]. Out of the 27 species identified, seven Peyssonneliales species and nine Corallinales species could not be assigned to recognized genera at this time. While two Corallinales species were assigned to the *Titanoderma*/*Lithophyllum* genera complex. Upon completion of BLAST searches, no sequences of described species or recognized genera returned a close match (< 97%) with these 16 species, therefore phylogenies were used to identify species to the lowest taxonomic level possible.

Our results also suggest that Lithophylloideae spp. 1–2 could be assigned to a new genus that includes taxa identified as *Titanoderma* but are paraphyletic to the clade that contains *Titanoderma pustulatum* [46,69,84]. Resolving and characterizing the identity of Lithophylloideae spp. 1–2 is warranted given the important ecological role of Lithophylloideae sp. 1 as a recruitment substrate for coral larvae.

The validity of the genus *Titanoderma* has been up for debate for decades. The original morphological distinction between *Titanoderma* and *Lithophyllum* is due to the basal layer of palisade cells and bistratose margins (*Titanoderma*) or basal layer of non-palisade cells and non-bistratose margins (*Lithophyllum*) [70, 85]. However, these diagnostic features can coexist within a single thallus of both *Lithophyllum* and *Titanoderma*, which resulted in the proposal of merging *Titanoderma* into *Lithophyllum* [51]. There is no consensus about this proposal because there is a predominance of different hypothallial cells (palisade cells and non-palisade cells) in both genera and the presence of a bistratose margin in *Titanoderma*. If *Titanoderma* is regarded as a distinct genus [46, 85] based on a phylogenetic delineation, Lithophylloideae sp. 3 and Lithophylloideae sp. 4 are *Titanoderma* species. Resolving the nomenclatural status of *Titanoderma* was beyond the scope of this study and therefore we opted to assign these species to the lowest taxonomic rank to which they could be positively identified (i.e., subfamily).

### Ecological significance of CCRA for *Acropora surculosa* recruitment

Despite their ability to recruit on a broad diversity of CCRA species, *Acropora* spp. have been reported to actively favor specific CCRA species as recruitment substrates when present. Prior to this study, it was unknown if and which CCRA species were preferred by *A. surculosa* as recruitment substrates. Here we report that *A. surculosa* larvae actively favor certain CCRA species over others, which supports similar findings for other *Acropora* species [10,34,36]. In other studies, *Titanoderma prototypum* (Foslie) Woelkerling, Y.M. Chamberlain & P.C. Silva has been characterized by low benthic cover, yet *Acropora palmata* and *Montastraea faveolata* significantly preferred this species over more abundant CCRA [35]. In other experimental studies in the Pacific [34], *T. prototypum* was the dominant of five CCRA species on coral recruitment tiles and also the favored CCRA species for recruitment and settlement. Despite *T. prototypum* abundance on the recruitment tiles, it was not commonly found in Moorea’s reef community [34]. The putative *Titanoderma* species in our study, i.e.

Lithophylloideae sp. 1, was also the dominant CCRA species on coral recruitment tiles. *Titanoderma* spp. are regarded to be a minor component of Pacific reefs (low benthic cover) and have been found in the mid and outer zones of reef systems [34,35,86]. The benthic cover of Lithophylloideae sp. 1 on Guam’s natural reefs has not been assessed and ongoing barcoding efforts of Guam’s CCRA have yet to detect the alga from natural reef communities. Based on the findings from this study, the presence of Lithophylloideae sp. 1 could be beneficial for reef recovery following disturbance events.

Lithophylloideae sp. 1 was morphologically distinct from the other 16 Corallinales species identified on the recruitment tiles. In white light, it had a deeper red, pink pigmentation with noticeably large conceptacles (Fig 1). Using a fluorescence setup (actinic exciter light and yellow barrier filter), Lithophylloideae sp. 1 displayed a fluorescent orange pigmentation that was not observed for the other 26 Corallinales species (Fig 1). This could signify that Lithophylloideae sp. 6 has a distinct chemical and microbial community that facilitates successful acroporid recruitment. It has been proposed that successful coral larval recruitment and settlement is largely due to the epiphytic microbiome communities and chemical signatures of CCRA [8, 32], which may be unique for a species [40]. *T. prototypum* is known to harbor distinct microbial community and metabolomic fingerprint compared to other CCRA [40, 87] in studies where *T. prototypum* was strongly favored for settlement and attachment by *Acropora cytherea* larvae.

Since adult coral colonies are sessile, competition for space is a major factor impacting the survival of a coral colony. Despite the species-richness reported for the recruitment tiles, only five CCRA species that were not Lithophylloideae members were associated with a coral recruit. *A. surculosa* larvae were able to recruit to Peyssonneliales sp. 4, *Polystrata* sp. 3, *Harveylithon* sp. 3, Lithophyllaceae sp. 4 and one Corallinales spp. Successful larval recruitment on non-favored CCRA species could in part be due to random recruitment and competition [88], however there are many abiotic and biotic factors that contribute to larval recruitment that are largely not understood. The ability for larvae to rank CCRA species for recruitment and their capability to recruit to non-favored CCRA species allows for coral colonies to successfully settle and reproduce in a variety of habitat types.

Ocean acidification has shown to alter the chemical cues to recognize Corallinales taxa by coral larvae and can lead to reduced recruitment [89]. Peyssonneliales taxa are believed to be less susceptible to ocean acidification [90] than the Corallinales [89]. The ability to recruit to Peyssonneliales taxa could be beneficial for coral larval recruitment in climate change scenarios that affect the growth of Corallinales taxa. *A. surculo*sa larvae recruited to two Peyssonneliales species (Peyssonneliales sp. 4 and *Polystrata* sp. 3). A study on *Goniastrea retiformis* in Guam [33] demonstrated that the larvae of this coral can also recruit to Peyssonneliales taxa although they are not the preferred recruitment substrates. Studies in which *T. prototypum* was the favored recruitment substrate [10], also found that coral recruit mortality was higher when growing on CCRA other than *T. prototypum.* This study did not examine the growth and survival of coral recruits in relation to CCRA substrates.

This study demonstrates that DNA sequencing can resolve a higher number of CCRA species in coral recruitment experiments compared to identifications that are only based on morphological features [79]. Given the high CCRA diversity on a small spatial scale, it is expected that the floristic diversity of CCRA in the tropical Pacific is severely understudied. Experiments that investigate ecological characteristics of different CCRA taxa are particularly valuable to assess and evaluate reef health in monitoring and environmental impact studies. This study identified that Acroporid larvae prefer to recruit on specific CCRA species, which can guide coral reef management and conservation programs. The favored Lithophylloideae sp. 1 has yet to be described and encountered on natural reef systems. Research on the role and diversity of CCRA in tropical reef communities is required to build a holistic view of reef ecosystem composition and ecological processes.

## Acknowledgements

We are grateful to Dr. Laurie Raymundo and the Raymundo Coral Lab for the use of the coral recruitment tiles for this research study. This manuscript benefitted from conversations and assistance regarding R scripting and data analysis by Andrew McInnis. A special thanks to Dr. Alex Kerr and Dr. Heroen Verbruggen for their invaluable time in extending their knowledge. MD, MM, and TS are indebted to the University of Guam for supporting studies that document and conserve the natural resources of Guam and the greater Micronesian region.

## Captions for Supplementary Materials

**S1 Fig. Seven-gene concatenated maximum likelihood phylogeny for the 17 Corallinales species from the recruitment tiles (boldface font) and reference sequences from Peña et al.** [69]. The tree was inferred by maximum likelihood with IQ-TREE [66]. Support values listed next to each node are ultrafast bootstrap approximations based on 1000 replicates. Scale bar represents nucleotide substitutions per site.

**S1 Table.** GenBank accession numbers for COI-5P, *psb*A, and *rbc*L sequences of specimens from the coral recruitment tiles, with indication of the order, taxon name, and herbarium accession number.

**S2 Table.** GenBank accession numbers of reference sequences from Peña et al. [69] used in the phylogenetic tree of Corallinales, S1 Fig.

**S3 Table.** GenBank accession numbers or BOLD Systems identification numbers (*) of COI-5P sequences of Peyssonneliales taxa used in Fig 3.

