## Supplemental Table 1 for "Community assessment of crustose calcifying red algae as coral recruitment substrates"

**S1 Table:** List of specimens from the coral recruitment tiles used to generate DNA sequences (COI-5P, psbA, rbcL) for this study, their Order, taxon name, Herbarium number, and corresponding GenBank accession numbers.

|  |  |  | GenBank accession numbers | | |
| --- | --- | --- | --- | --- | --- |
| Order | **Taxon** | **Guam Herbarium** | **COI-5P** | ***psb*A** | ***rbc*L** |
| Corallinales |  |  |  |  |  |
|  | *Harveylithon* sp. 1 | GH0015508 | MZ824140 | MZ824164 | MZ824213 |
|  | *Harveylithon* sp. 2 | GH0015432 | MZ824109 | MZ824144 | MZ824198 |
|  | *Harveylithon* sp. 3 | GH0015497 | MZ824133 | MZ824162 |  |
|  | *Hydrolithon* sp. 1 | GH0015464 | MZ824118 | MZ824154 | MZ824204 |
|  | Lithophyllaceae sp. 1 | GH0015475 | MZ824122 | MZ824157 | MZ824207 |
|  | Lithophyllaceae sp. 2 | GH0015504 | MZ824137 |  | MZ824212 |
|  | Lithophyllaceae sp. 3 | GH0015456 |  | MZ824151 | MZ824201 |
|  | Lithophyllaceae sp. 4 | GH0015479 |  | MZ824158 | MZ824208 |
|  | Lithophyllaceae sp. 5 | GH0015458 |  | MZ824152 | MZ824202 |
|  | Lithophyllaceae sp. 6 | GH0015496 |  | MZ824161 | MZ824210 |
|  | Lithophyllaceae sp. 7 | GH0015467 |  | MZ824155 |  |
|  | Lithophylloideae sp. 1 | GH0015421 | MZ824103 |  |  |
|  | Lithophylloideae sp. 1 | GH0015425 | MZ824104 |  |  |
|  | Lithophylloideae sp. 1 | GH0015426 | MZ824105 |  |  |
|  | Lithophylloideae sp. 1 | GH0015427 | MZ824106 |  |  |
|  | Lithophylloideae sp. 1 | GH0015428 | MZ824107 |  |  |
|  | Lithophylloideae sp. 1 | GH0015429 | MZ824108 |  |  |
|  | Lithophylloideae sp. 1 | GH0015441 | MZ824145 |  |  |
|  | Lithophylloideae sp. 1 | GH0015446 | MZ824111 |  |  |
|  | Lithophylloideae sp. 1 | GH0015447 | MZ824112 | MZ824147 |  |
|  | Lithophylloideae sp. 1 | GH0015449 | MZ824113 | MZ824148 |  |
|  | Lithophylloideae sp. 1 | GH0015450 | MZ824114 | MZ824149 | MZ824200 |
|  | Lithophylloideae sp. 1 | GH0015453 | MZ824115 |  |  |
|  | Lithophylloideae sp. 1 | GH0015455 | MZ824150 |  |  |
|  | Lithophylloideae sp. 1 | GH0015460 | MZ824116 |  |  |
|  | Lithophylloideae sp. 1 | GH0015461 | MZ824117 |  |  |
|  | Lithophylloideae sp. 1 | GH0015469 | MZ824119 |  |  |
|  | Lithophylloideae sp. 1 | GH0015473 | MZ824121 |  | MZ824206 |
|  | Lithophylloideae sp. 1 | GH0015476 | MZ824123 |  |  |
|  | Lithophylloideae sp. 1 | GH0015477 | MZ824124 |  |  |
|  | Lithophylloideae sp. 1 | GH0015481 | MZ824125 |  |  |
|  | Lithophylloideae sp. 1 | GH0015482 | MZ824126 |  |  |
|  | Lithophylloideae sp. 1 | GH0015484 | MZ824127 |  |  |
|  | Lithophylloideae sp. 1 | GH0015485 | MZ824128 |  |  |
|  | Lithophylloideae sp. 1 | GH0015490 | MZ824130 |  |  |
|  | Lithophylloideae sp. 1 | GH0015494 | MZ824131 |  |  |
|  | Lithophylloideae sp. 1 | GH0015501 | MZ824135 |  |  |
|  | Lithophylloideae sp. 1 | GH0015502 | MZ824136 |  |  |
|  | Lithophylloideae sp. 1 | GH0015506 | MZ824138 |  |  |
|  | Lithophylloideae sp. 1 | GH0015507 | MZ824139 |  |  |
|  | Lithophylloideae sp. 1 | GH0015509 | MZ824141 |  |  |
|  | Lithophylloideae sp. 1 | GH0015510 | MZ824142 |  |  |
|  | Lithophylloideae sp. 2 | GH0015495 | MZ824132 | MZ824160 |  |
|  | Lithophylloideae sp. 3 | GH0015470 | MZ824120 | MZ824156 | MZ824205 |
|  | Lithophylloideae sp. 3 | GH0015512 | MZ824143 |  | MZ824214 |
|  | Lithophylloideae sp. 4 | GH0015462 |  | MZ824153 | MZ824203 |
|  | Lithophylloideae sp. 4 | GH0015498 | MZ824134 | MZ824163 | MZ824211 |
|  | Lithophylloideae sp. 4 | GH0015511 |  | MZ824165 |  |
|  | *Neogoniolithon* sp. 1 | GH0015443 | MZ824110 | MZ824146 | MZ824199 |
|  | *Porolithon* sp. 1 | GH0015486 | MZ824129 | MZ824159 | MZ824209 |
| Peyssonneliales |  |  |  |  |  |
|  | Peyssonneliales sp. 1 | GH0015503 | MZ824195 |  |  |
|  | Peyssonneliales sp. 1 | GH0015451 | MZ824180 |  |  |
|  | Peyssonneliales sp. 1 | GH0015487 | MZ824191 |  |  |
|  | Peyssonneliales sp. 2 | GH0015430 | MZ824168 |  |  |
|  | Peyssonneliales sp. 2 | GH0015452 | MZ824181 |  |  |
|  | Peyssonneliales sp. 3 | GH0015471 | MZ824187 |  |  |
|  | Peyssonneliales sp. 3 | GH0015474 | MZ824188 |  |  |
|  | Peyssonneliales sp. 4 | GH0015422 | MZ824166 |  |  |
|  | Peyssonneliales sp. 4 | GH0015434 | MZ824171 |  |  |
|  | Peyssonneliales sp. 4 | GH0015438 | MZ824173 |  |  |
|  | Peyssonneliales sp. 4 | GH0015442 | MZ824176 |  |  |
|  | Peyssonneliales sp. 4 | GH0015444 | MZ824177 |  |  |
|  | Peyssonneliales sp. 4 | GH0015445 | MZ824178 |  |  |
|  | Peyssonneliales sp. 4 | GH0015465 | MZ824184 |  |  |
|  | Peyssonneliales sp. 4 | GH0015483 | MZ824190 |  |  |
|  | Peyssonneliales sp. 4 | GH0015488 | MZ824192 |  |  |
|  | Peyssonneliales sp. 4 | GH0015499 | MZ824193 |  |  |
|  | Peyssonneliales sp. 5 | GH0015440 | MZ824175 |  |  |
|  | Peyssonneliales sp. 6 | GH0015423 | MZ824167 |  |  |
|  | Peyssonneliales sp. 6 | GH0015431 | MZ824169 |  |  |
|  | Peyssonneliales sp. 6 | GH0015468 | MZ824186 |  | - |
|  | Peyssonneliales sp. 6 | GH0015505 | MZ824196 |  |  |
|  | Peyssonneliales sp. 6 | GH0015513 | MZ824197 |  |  |
|  | Peyssonneliales sp. 7 | GH0015433 | MZ824170 |  |  |
|  | Peyssonneliales sp. 7 | GH0015448 | MZ824179 |  |  |
|  | Peyssonneliales sp. 7 | GH0015454 | MZ824182 |  |  |
|  | *Polystrata* sp. 1 | GH0015459 | MZ824183 |  |  |
|  | *Polystrata* sp. 2 | GH0015436 | MZ824172 |  |  |
|  | *Polystrata* sp. 3 | GH0015439 | MZ824174 |  |  |
|  | *Polystrata* sp. 3 | GH0015466 | MZ824185 |  |  |
|  | *Polystrata* sp. 3 | GH0015480 | MZ824189 |  |  |
|  | *Polystrata* sp. 3 | GH0015500 | MZ824194 |  |  |
