## Supplemental Table 2 for "Community assessment of crustose calcifying red algae as coral recruitment substrates"

**S2 Table.** List of taxa for Corallinales species delimitation included in the phylogenetic analyses detailing GenBank accession number for each marker.

|  |  |  | GenBank accession numbers | | | | | | |
| --- | --- | --- | --- | --- | --- | --- | --- | --- | --- |
| Taxon | **Order** | **Species** | **COI** | ***psbA*** | ***rbc*L** | **23S rRNA** | **SSU rRNA** | **LSU rRNA** | **EF2** |
| Taxon 001 | Corallinales | Lithophyllaceae sp. (voucher ID MAD2358, Madagascar) | **MT325717** | **MT325762** |  |  |  | **MT325700** | - |
| Taxon 001a | Corallinales | Lithophyllaceae sp. (voucher ID VPF00502A, Atlantic Spain) | **MT325711** | **MT325756** | **MT325734** |  |  | **MT325687** |  |
| Taxon 002 | Corallinales | *Metagoniolithon radiatum* |  | GQ917496 |  |  | GQ917432 | GQ917369 |  |
| Taxon 002b | Corallinales | *Metagoniolithon stelliferum* |  | GQ917497 |  |  | GQ917433 | GQ917370 |  |
| Taxon 003 | Corallinales | Lithophyllaceae sp.(voucher ID MAD1024B, Madagascar) | **MT325705** | **MT325748** |  |  |  | **MT325678** |  |
| Taxon 003a | Corallinales | Lithophyllaceae sp. (voucher ID MAD938B, Madagascar) | **MT325709** | **MT325754** |  |  | **MT325665** | **MT325685** |  |
| Taxon 004 | Corallinales | *Dawsoniolithon* sp. (as Uncultured Corallinales, Bittner et al. 2011) | GQ917265 | GQ917454 |  |  | GQ917399 | GQ917328 |  |
| Taxon 005 | Corallinales | *Dawsoniolithon conicum* (as *Pneophyllum conicum*, Bittner et al. 2011) | GQ917272 | GQ917464 |  | HQ423076 | GQ917408 | GQ917337 |  |
| Taxon 006 | Corallinales | *Harveylithon* sp. (as Uncultured Corallinales, voucher ID LBC0600, Bittner et al. 2011) | GQ917264 | GQ917453 | **MT325719** |  | GQ917398 | GQ917327 |  |
| Taxon 007 | Corallinales | *Harveylithon* sp. (voucher ID MAS304_1C, Oman) | **MT325714** | **MT325758** |  |  |  | **MT325693** |  |
| Taxon 008 | Corallinales | *Porolithon* sp. (as *Hydrolithon onkodes*, voucher ID LBC0820_P3, Bittner et al. 2011) | GQ917291 | GQ917483 | **MT325738** | KM073308 | GQ917373 | GQ917357 |  |
| Taxon 009 | Corallinales | *Porolithon* sp. (as *Hydrolithon* sp., Bittner et al. 2011) | GQ917286 | GQ917478 |  | HQ42271 | GQ917422 | GQ917352 |  |
| Taxon 010 | Corallinales | *Porolithon* *onkodes* (as *Hydrolithon* sp., voucher ID LBC0678, Bittner et al. 2011) | GQ917276 | GQ917469 |  |  | GQ917412 | GQ917341 |  |
| Taxon 011 | Corallinales | *Porolithon* sp. (as *Hydrolithon* sp., Bittner et al. 2011) | GQ917287 | GQ917479 |  |  | GQ917423 | GQ917353 |  |
| Taxon 012 | Corallinales | *Porolithon* sp.(as *Hydrolithon* sp., voucher ID LBC0882, Bittner et al. 2011) | GQ917301 | GQ917493 | **MT325720** |  | GQ917429 | GQ917366 |  |
| Taxon 013 | Corallinales | Lithophyllaceae sp. (voucher ID MAD2344, Madagascar) | **MT325715** | **MT325759** | **MT325737** |  |  | **MT325694** |  |
| Taxon 014 | Corallinales | Lithophyllaceae sp. (voucher ID NVT1054B, Vietnam) | **MT325704** | **MT325747** | **MT325722** |  | **MT325662** | **MT325677** |  |
| Taxon 015 | Corallinales | *Adeylithon bosencei* (as Hydrolithon sp., voucher ID LBC0720, Bittner et al. 2011) | GQ917284 | GQ917476 | **MT325731** |  | GQ917420 | GQ917349 |  |
| Taxon 016 | Corallinales | *Hydrolithon* cf. *boergesenii* | GQ917257 | GQ917447 |  |  | GQ917378 | GQ917321 |  |
| Taxon 017 | Corallinales | *Hydrolithon reinboldii* | GQ917293 | GQ917485 |  | HQ423071 | GQ917376 | GQ917359 |  |
| Taxon 019 | Corallinales | *Chamberlainium* sp. (voucher ID LLG4403B, Australia) |  | **MT325753** |  |  |  | **MT325684** |  |
| Taxon 020 | Corallinales | *Spongites hyperellus* |  | GQ917495 |  |  | GQ917431 | GQ917368 |  |
| Taxon 021 | Corallinales | *Pneophyllum fragile* |  | KT783426 |  |  |  |  |  |
| Taxon 022 | Corallinales | *Pneophyllum cetinaensis* (voucher ID PC0145164, Žuljević et al. 2016) |  | KT783433 |  |  |  | **MT325690** |  |
| Taxon 023 | Corallinales | *Spongites yendoi* |  | DQ167907 | KT184848 |  | EF628234 |  |  |
| Taxon 024 | Corallinales | Lithophyllaceae sp. (as Uncultured Corallinales, voucher ID LBC0707, Bittner et al. 2011) | GQ917589 | GQ917960 | **MT325725** |  | **MT325664** | **MT325682** |  |
| Taxon 025 | Corallinales | *Lithophyllum* sp. (voucher LBC0714, Bittner et al. 2011) | GQ917282 | GQ917474 | **MT325735** |  | GQ917418 | GQ917347 |  |
| Taxon 026 | Corallinales | *Lithophyllum hibernicum* | GQ917250 | GQ917440 | KR708590 | KM073316 | GQ917385 | GQ917313 |  |
| Taxon 027 | Corallinales | *Lithophyllum byssoides* (voucher ID VPF00306, Atlantic Spain) | **MT325702** | **MT325743** | KR708574 |  | JQ896251 | **MT325673** |  |
| Taxon 028 | Corallinales | *Lithophyllum* sp. (voucher ID MAD0081, Madagascar) | **MT325707** | **MT325750** |  |  | JQ896235 | **MT325680** |  |
| Taxon 029 | Corallinales | *Lithophyllum* sp. (voucher ID LBC0946, Bittner et al. 2011) | GQ917691 | GQ918126 | **MT325732** | HQ421552 | **MT325667** | HQ422479 |  |
| Taxon 031 | Corallinales | *Tinanoderma* sp. | KJ418416 | KJ418413 | KJ652016 |  |  | KJ412335 |  |
| Taxon 032 | Corallinales | *Lithophyllum pustulatum* (voucher ID VPF00095, Atl. Spain) | **MT325701** | **MT325742** | KM369168 | KM073327 | KM073290 | **MT325672** |  |
| Taxon 033 | Corallinales | *Lithothrix aspergillum* | JQ615866 | JQ422237 | HQ322336 |  |  |  | JQ422275 |
| Taxon 034 | Corallinales | *Lithophyllum kotschyanum* |  | AB576029 | KX020467 | KM073321 | AB576008 | KM977984 |  |
| Taxon 035 | Corallinales | *Tinanoderma* sp. | GQ917285 | GQ917477 |  |  | GQ917421 | GQ917350 |  |
| Taxon 036 | Corallinales | *Lithophyllum* sp. (voucher ID LBC0680, Bittner et al. 2011) | GQ917277 | GQ917470 | **MT325723** |  | GQ917413 | GQ917342 |  |
| Taxon 037 | Corallinales | *Amphiroa* sp. (voucher ID LBC0865, Bittner et al. 2011) | GQ917299 | GQ917491 | **MT325726** |  | GQ917428 | GQ917364 |  |
| Taxon 038 | Corallinales | *Amphiroa* sp. | GQ917246 | GQ917435 |  |  | GQ917380 | GQ917308 |  |
| Taxon 039 | Corallinales | *Amphiroa* sp. (voucher ID LBC0708, Bittner et al. 2011) | GQ917280 | GQ917472 | **MT325728** |  | GQ917416 | GQ917345 |  |
| Taxon 040 | Corallinales | *Amphiroa fragilissima* | GQ917303 | GQ917498 | U04039 | KM044012 | KY987580 | EF033599 |  |
| Taxon 054 | Corallinales | *Mastophora/Lithoporella* | GQ917260 | GQ917449 |  |  | GQ917394 | GQ917323 |  |
| Taxon 055 | Corallinales | *Jania longifurca* | GQ917251 | GQ917441 | KM369140 |  | GQ917386 | GQ917314 |  |
| Taxon 056 | Corallinales | *Jania* sp. | GQ917514 | GQ917712 |  | HQ421368 |  | HQ422166 |  |
| Taxon 057 | Corallinales | *Jania rubens* (voucher ID VPF00439, Atlantic Spain) | **MT325713** | MK308537 | KM044024 | KM044014 | KM044029 | **MT325691** |  |
| Taxon 058 | Corallinales | *Jania sagittata* | JQ615844 | JQ422232 | KC134331 |  | KM369032 | KC157591 | KC130175 |
| Taxon 069 | Corallinales | *Mastophora rosea* (voucher ID LBC0866, Bittner et al. 2011) | GQ917300 | GQ917492 | **MT325729** |  | **MT325666** | GQ917365 |  |
| Taxon 070 | Corallinales | *Mastophora pacifica* | GQ917302 | GQ917494 | KM369152 | HQ420920 | GQ917430 | GQ917367 |  |
| Taxon 081 | Corallinales | *Spongites* sp. (voucher ID LLG2579, Rösler et al. 2016) | KP682496 | **MT325752** |  |  |  | **MT325683** |  |
| Taxon 082 | Corallinales | *Spongites fruticulosus* (voucher ID VPF00027, Rösler et al. 2016) | **MT325710** | **MT325755** |  | KM073335 | KM073306 | **MT325686** |  |
| Taxon 083 | Corallinales | *Arthrocardia corymbosa* |  | JQ917408 | JN701475 | HQ421500 |  |  |  |
| Taxon 084 | Corallinales | *Chiharaea bodegensis* | HM918942 | JQ677011 | JQ677000 |  | KC157576 | KC157588 | KC130170 |
| Taxon 085 | Corallinales | *Calliarthron cheilosporioides* | JQ615594 | JQ422199 | HQ322299 |  | CTU60944 |  | JQ422270 |
| Taxon 086 | Corallinales | *Ellisolandia elongata* | JQ615843 | JQ422231 | JX315327 |  | FM180099 |  | JQ422258 |
| Taxon 087 | Corallinales | *Corallina caespitosa* | GQ917248 | GQ917438 | JQ615683 | KC478072 | GQ917383 | GQ917311 | KC130168 |
| Taxon 088 | Corallinales | *Neogoniolithon* sp. (voucher ID FRA1402, Guadeloupe, West Indies) | KP682495 | KP682501 |  |  | **MT325668** | **MT325692** |  |
| Taxon 089 | Corallinales | *Neogoniolithon* sp. (voucher ID LBC0584, Bittner et al. 2011) | GQ917262 | GQ917451 | **MT325730** |  | GQ917396 | GQ917325 |  |
| Taxon 090 | Corallinales | *Neogoniolithon* sp. | GQ917290 | GQ917482 |  |  | GQ917424 | GQ917356 |  |
| Taxon 091 | Corallinales | *Neogoniolithon* sp. (voucher ID LBC0843, Bittner et al. 2011) | GQ917297 | GQ917489 | **MT325727** |  | GQ917434 | GQ917362 |  |
| Taxon 092 | Corallinales | *Neogoniolithon brassica-florida* (voucher ID VPF00284, Med. France) | KM392368 | **MT325745** |  | HQ422477 | JQ896257 | **MT325675** |  |
| Taxon 093 | Corallinales | *Neogoniolithon* sp. (voucher ID LBC0433, Bittner et al. 2011) | GQ917253 | GQ917442 | **MT325733** |  | GQ917388 | GQ917316 |  |
| Taxon 094 | Corallinales | *Neogoniolithon* sp. | GQ917274 | GQ917466 |  |  | GQ917410 | GQ917339 |  |
| Taxon 095 | Corallinales | *Amphiroa valonioides* | HQ422698 |  |  | HQ421023 |  | HQ422411 |  |
| Taxon 096 | Corallinales | *Amphiroa foliacea* | HQ422626 |  |  | HQ420962 |  | HQ421910 |  |
| Taxon 097 | Corallinales | *Lithophyllum* sp. | HQ422960 |  |  |  |  | HQ422022 |  |
| Taxon 098 | Corallinales | *Lithophyllum insipidum* | HQ423068 |  |  | HQ421545 |  | HQ422473 |  |
| Taxon 102 | Corallinales | *Jania* sp. | HQ423038 |  |  | HQ421370 |  | HQ422269 |  |
| Taxon 103 | Corallinales | *Jania* sp. | HQ422629 |  |  | HQ421458 |  | HQ421768 |  |
| Taxon 106 | Corallinales | *Spongites* sp. | HQ422715 |  |  | HQ420977 |  | HQ421807 |  |
| Taxon 107 | Corallinales | *Porolithon gardineri* | HQ423069 |  |  | HQ421546 |  | HQ422474 |  |
| Taxon_131 | Corallinales | *Pneophyllum coronatum* |  | DQ168008 |  |  | KM369048 |  |  |
| Taxon_132 | Corallinales | *Pneophyllum fragile* |  | FJ361387 | KM369155 |  | KM369043 |  |  |
| Taxon_133 | Corallinales | *Metamastophora flabellata* |  |  |  |  | AY234240 |  |  |
| Taxon_134 | Corallinales | *Metamastophora flabellata* |  |  |  |  | AY234239 |  |  |
| Taxon_135 | Corallinales | *Mastophora* sp. |  |  |  |  | MG826386 | MG821596 |  |
| Taxon_136 | Corallinales | *Parvicellularium leonardii* |  | MG851066 |  |  | MG826383 | MG821592 |  |
| Taxon_137 | Corallinales | *Parvicellularium* sp. |  | MG851074 |  |  |  | MG821601 |  |
| Taxon_138 | Corallinales | *Porolithon* cf. *craspedium* |  | MG851057 |  |  | MG826374 | MG821582 |  |
| Taxon_139 | Corallinales | *Spongites* sp. | MG851045 | MG851062 |  |  | MG826379 | MG821588 |  |
| Taxon_140 | Corallinales | *Spongites* sp. | MG851047 | MG851073 |  |  | MG826390 | MG821600 |  |
| Taxon_141 | Corallinales | *Neogoniolithon* sp. | MG851044 | MG851061 |  |  | MG826378 | MG821586 |  |
| Taxon_142 | Corallinales | *Chamberlainium* sp. |  | MG851078 |  |  | MG826394 |  |  |
| Taxon_143 | Corallinales | *Harveylithon munitum* |  | KM407531 |  |  |  | KM073336 |  |
| Taxon_144 | Corallinales | *Harveylithon rupestre* |  | KM407535 |  |  | KM073303 |  |  |
