## Supplemental Table 3 for "Community assessment of crustose calcifying red algae as coral recruitment substrates"

**S3 Table. List of Peyssonneliales taxa for species delimitation included in the phylogenetic analyses detailing GenBank accession number for COI.** * represent BOLD system identification number.

|  |  | **GenBank accession numbers** |
| --- | --- | --- |
| **Taxa** | **Specimen Voucher** | **COI** |
| *Ramicrusta nanhaiensis* | GWS002520 | JX969713 |
| *Polystrata* cf. *dura* | VT097 | ABMMC1983-07* |
| *Polystrata* cf. *dura* | VT159 | ABMMC1984-07* |
| *Metapeyssonnelia* sp. | 6552 | ABMMC320-06* |
| *Metapeyssonnelia* sp. | 6558 | ABMMC323-06* |
| *Metapeyssonnelia* sp. | VT086 | ABMMC1862-07* |
| *Riquetophycus polypus* | HSY-2014a | KJ398161 |
| *Sonderophycus fervens* | VT061 | JX969700 |
| *Cruoriella* sp. | VT166 | ABMMC1861-07* |
| *Peyssonnelia squamaria* | GWS018179 | JX969741 |
| *Incendia crenata* | VT095 | JX969714 |
| *Bonnemaisonia asparagoides* | GWS040634 | MN184297 |
