## Supplementary figures and images for "Community assessment of crustose calcifying red algae as coral recruitment substrates"

### Supplemental Figure 1

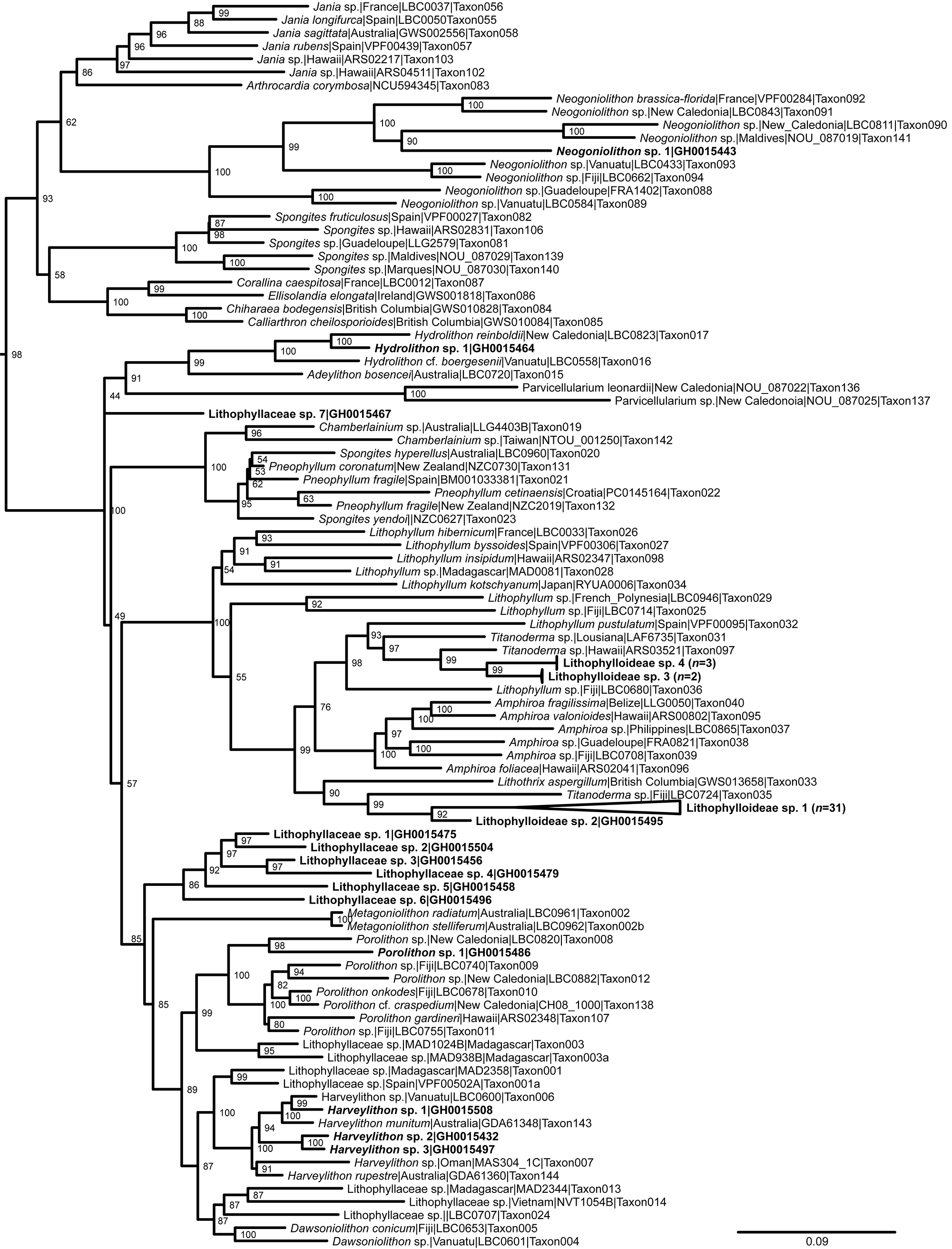
